# Upscaling of Microbial Electrolysis Cell Integrating Microbial Electrosynthesis: Insights, Challenges and Perspectives

**DOI:** 10.1101/609909

**Authors:** Jiang-Hao Tian, Rémy Lacroix, Elie Desmond-Le Quéméner, Chrystelle Bureau, Cédric Midoux, Théodore Bouchez

## Abstract

Recent development of microbial electrochemical technologies has allowed microbial electrosynthesis (MES) of organic molecules with microbial electrolysis cell treating waste organic matter. An electrolytic cell with a MES cathode (ME-ME cell) can produce soluble organic molecules with higher market price than biomethane, and thus satisfy both economic and environmental interest. However, the sustainability of bioanode activity could become a major concern. In this work, a 15-liter ME-ME reactor was designed with specific electrode configurations. An electrochemical model was established to assess the feasibility and possible performance of the design, considering the “aging” effect of the bioanode. The reactor was then built and operated for performance evaluation as well as bioanode regeneration assay. Biowaste from an industrial deconditioning platform was used as substrate for bioanode. The COD removal rate in the anodic chamber reached 0.83 g day^-1^ L^-1^ of anolyte and the anodic coulombic efficiency reached 98.6%. Acetate was produced with a rate of 0.53 g day^-1^ L^-1^ of catholyte, reaching a maximum concentration of 8.3 g L^-1^. A potential difference was applied between the bioanode and biocathode independent of reference electrodes. The active biocathode was dominated by members of the Genus *Pseudomonas*, rarely reported so far for MES activity.

## 1 Introduction

The environmental biorefinery consists of transformation of organic wastes into resources [1]. According to this concept, waste generated by our society is recycled and reused in processes in which they are transformed into value-added products instead of being directly mineralized and/or discharged in nature [2]. Anaerobic digestion (AD) is a simple, spontaneous and robust process and a well-established example of a technology able to recover organic waste into methane. In Europe, 1.35×10^7^ tons of biogas was produced in 2014, corresponding to 3% of natural gas consumption of the continent [3]. It is however noteworthy that the economic viability of AD partly depends on government grants in many European countries because biomethane is not yet a competitive energy source on the market [4]. Will it become competitive in the future? According to Bioenergy Europe (formerly known as AEBIOM), the technical potential of biomethane in EU is about 78 billion Nm^3^, corresponding to 15.6% of the 500 billion Nm^3^ annual natural gas consumption of Europe [5, 6]. In other words, an exhaustive exploration of EU’s biomethane potential would merely fulfill less than 1/6 of the continent’s natural gas requirement. Therefore, as an energy product, biomethane will always face severe competition from other energy sources, probably more cost-effective. Transforming the available organic matters into products of higher value may thus be a more interesting approach. Some biological processes such as fermentation and microbial electrochemical technologies (MET) can produce carboxylic acids and alcohols from residual organic matters. Precursors of various chemical processes, these platform molecules have considerable potential to improve the profitability of environmental biorefinery industries [7].

Intensive researches have been devoted to production of chemicals from organic wastes at laboratory scale, resulting in a plethora of possibilities [8, 9, 10]. Commonly used fermentation processes can produce H_2_ (sometimes with methane) and different types of soluble organic molecules [11]. Gasification-fermentation process consists of a syngas production step then a gas fermentation step in which CO_2_ and CO are transformed into acetyl-CoA though the Wood-Ljungdahl pathway catalyzed by acetogens, using H2 as electron donor. This energy consuming process permits the valorization of recalcitrant lignocellulosic wastes [12]. The recent development of METs has created new application possibilities by combining biological and electrochemical processes to generate electricity or other products of interest. Electro-fermentation uses a polarized electrode to influence both the fermentative microbiota and their metabolism [11], allowing high-rate cathodic production of H_2_ and 1,3-propandiol from glucose and glycerol, respectively [13, 14]. The major challenge faced by these fermentation processes is the difficulties in purification of waste-derived products, given the fact that the separation of low-carbon acids and alcohols remains unsatisfactory with current technologies [15].

There are other non-fermentative METs. The concept of microbial fuel cells (MFCs) for electricity production from organic matter dates back almost 100 years [16]. A MFC can be converted to a microbial desalination cell (MDC) able to remove salts from seawater by inserting a pair of anion exchange membrane (AEM) and cation exchange membrane (CEM) between the anodic and the cathodic chamber [17]. Microbial electrolysis cells (MECs) were firstly proposed by two independent groups [18, 19] with the intention of producing reduced molecules (majorly H_2_) at the cathode by biologically oxidizing organic matter at the anode [20]. In addition to their main purposes, MFCs and MECs are also capable of removing recalcitrant pollutants or recovering valuable metals, with the help of specific anode or cathode potential [21]. Microbial electrosynthesis (MES) was discovered in 2010, opening new possibilities of producing soluble multi-carbons compounds from renewable electricity and CO_2_ with high selectivity, high rate and with high energy efficiency [22]. Although the mechanism of MES remains unclear, some evidences suggested that it could be highly H_2_ dependent, implying the involvement of the Wood-Ljungdahl pathway [23]. This cathodic process can produce a wide range of green chemistry precursors including fatty acids (acetate, butyrate or caporate) or alcohols (methanol, ethanol) using either defined cultures or consortia [24]. With a suitable community, acetate could be accumulated at high concentrations within a short period of time with close to 100% of e^-^ recovery [25].

Conventional MES systems usually use abiotic anodes made of expensive metals that are able to tolerate high voltages (3-5 V) [26, 27, 28]. This configuration requires high investment cost and high operation cost due to electricity consumption which are undesirable for large scale facilities [29]. The idea of using waste oxidation at a bioanode (equivalent to a MEC half-cell) to drive a MES biocathode has been mentioned by several authors since 2010 [30, 31, 32]. Recently, Desmond-Le Quéméner et al. presented the first laboratory scale proof of MEC fueled MES (ME-ME) cell (under review). In addition to the four-fold reduction in electric power consumption, this configuration included a CEM allowing a physical separation of the contaminated waste stream in the anodic chamber and the product molecule in the cathodic chamber, which is an important advantage for commercialization of the product molecules.

It is one thing to prove a concept in the laboratory, but quite another to implement the same technology at a large scale. The number of actual MET pilot projects is still very limited compared to lab-scale experiments because the list of unsolved up-scaling issues is remarkable [33]. Large varieties of parameters including configuration, materials, dimension, electrodes, microbial community, temperature, pH, substrate flow and cost-effectiveness critically govern the performance of a microbial electrochemical system. Pilot design and operation are thus a time-consuming and risky work: any single issue could lead to process failure [34, 35]. That may explain the lack of publication in this domain. It is however noteworthy that pilot assays are maybe the only way to apply acquired knowledge on a larger scale and reveal non-considered issues about up scaling.

Based on the previous work of Desmond-Le Quéméner et al., the MEC fueled MES technology (ME-ME cell) seems to be a competitive candidate for future environmental biorefinery, although many fundamental and practical issues remain to be addressed. We’d like to highlight two of them here. Firstly, in industrial environments, use of reference electrodes entrails numerous problems such as operation cost, implementation limits, breakage risks and imprecision due to fouling. It is thus interesting to have a system operating with a potential difference, given the fact that both the anode and cathode potential should be kept in a certain range. Secondly, in a MEC the anodic biocatalysts require activation and maintenance. For the bioanode, initial inoculum and start-up strategies have been disucussed by many authors [36]. However, we found very little data about bioanode activity maintenance and regeneration. Indeed, during long time operations, the anodic microbiota could gradually lose the desired electroactivity, in spite of sufficient substrate supply and adapted potential. We have observed this “aging” effect of bioanodes many times on different reactors. It is evidently a major obstacle to scaling up MECs. Frequently exchange or restart the bioanode would restore the activity of bioanodes but is clearly too slow and too costly for large-scale operations. It is thus indispensable to develop simple and efficient regeneration methods [37].

In this work, we designed a 15 L reactor integrating a double-plate bioanode and a granular carbon bed MES biocathode, operated completely with a potential difference and equipped with an extraction compartment. The reasons underlying this specific design are explained and discussed further in this article. The effect of the reactor’s bioanode “aging” was analyzed using a bioelectrochemical model of the reactor. The reactor was then constructed implemented and operated for over one year using industrial biowaste as anodic substrate. The long-term operation proved the feasibility of the technology but also allowed to identify multiple issues that can cause a decrease of reactor performances over time. We especially present and discuss two methods aiming at regenerating the bioelectrochemical activity of aged bioanodes.

## 2 Materials and Methods

### 2.1 Reactor Design

A filter-press type reactor named “TRL4” was designed in collaboration with 6TMIC, France (Patent PN001017). The main material used was the poly(methyl methacrylate) (PMMA). Briefly, it consists three main parts as showed in Figure 1:

**Figure 1.**
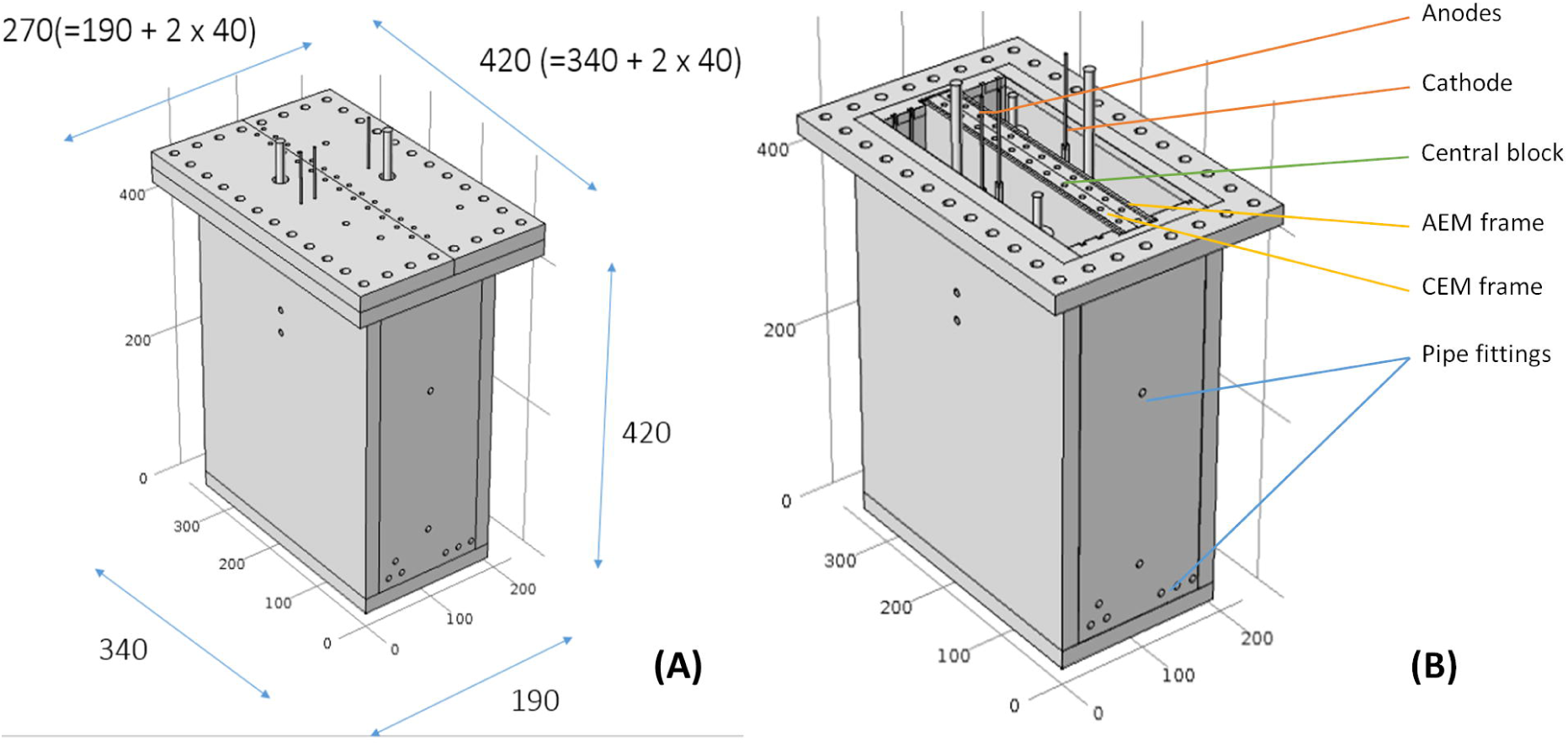
Overview 3D plans of the TRL4 reactor with (A) showing the reactor size with lids, (B) showing the overall configuration.

(i) The reactor body is a 340×190×420 (exterior) mm container with fixed walls (15 mm thickness) and bottom (20 mm thickness). Pipefittings for liquid circulation and gas injection were fixed on the walls;

(ii) An AEM and a CEM (Fumatech, Germany) frame with pasted gaskets and a hollow central block are disposed inside the reactor body to separate the inner space into three chambers: the anodic chamber (**AC**) (5.25L), the inter-membrane chamber (**IC**) (the space in the central block) (2.5L) and the cathodic chamber (**CC**) (5.25L);

(iii) The lid of AC and the lid of the CC can be fixed with screws at the top the reactor body and the central block, with a thick gasket between them ensuring gas tightness.

Electrode structure is important for the performance of METs. For the anode, Blanchet, et al. [38] showed that the electroactive biofilm formed only on the electrode surface of biowaste-fed bioanodes. On the other hand, surface cleaning could be a simple but efficient method for bioanode activity regeneration because a clean electrode surface is favorable to the colonization of members of the Genus *Geobacter*. For these reasons, anode of the TRL4 reactor consisted of two removable parallel flat plates, which allows to clean one of the anode plate while the other remains in function. As for the material, carbon felt is a widely used bioanode support that is friendly to biofilm but lacks rigidity and shear strength. Meanwhile, stainless steel is a promising bioanode material according to Pocaznoi, et al. [39]. Therefore, anode plates were made of 316L stainless steel frame and six carbon tissue strips (260X20mm) (Paxitech, France) that were fixed between two stainless steel grills (Goodfellow, UK), so that the stripes were protected and electrolyte could circulate freely through the anode plates (Figure 2). On the cathode, hydrogen produced by water electrolysis is an essential mediator in the MES [23]. On the other hand, no clear evidence showed that MES bacteria are surface-attached. For these reasons, a massive carbon granular cathode was chosen to maximize the hydrogen-producing surface, which could improve H_2_ uptake by MES bacteria such as homoacetogenic species. Four stainless steel baskets were filled with a total mass of 1.2 kg of carbon granules (SGL carbon, Germany) and were fixed on the frame. Each electrode frame had a fixed 2 mm diameter stainless steel rod as current collector.

**Figure 2.**
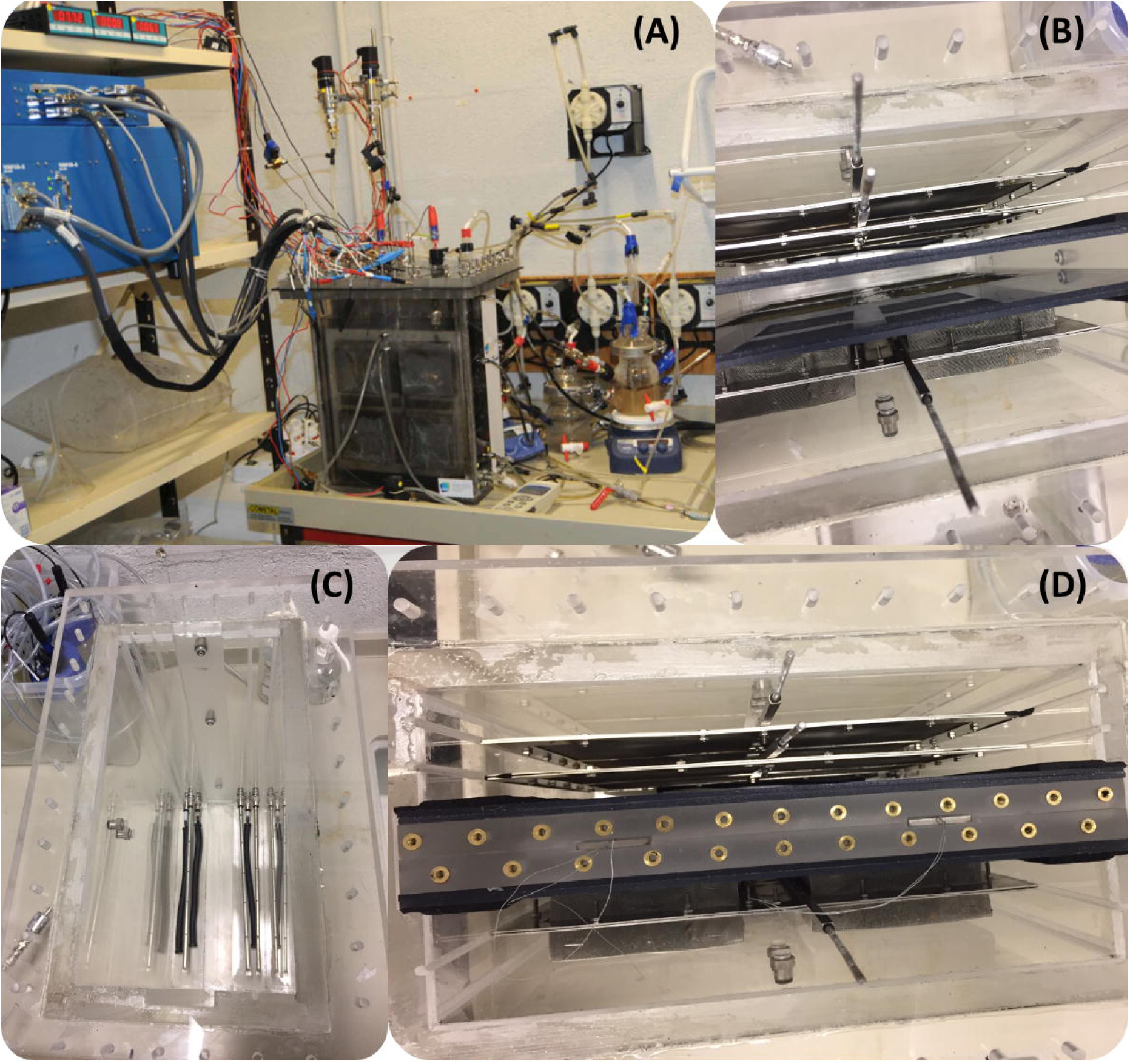
The integral system (A), the reactor body with electrodes and membrane frames (B), the reactor body with liquid injection pipes and gas diffusers (C) and the reactor body with electrodes, membrane frames and the center block (D).

Each reactor chamber was connected in a recirculation system including a continuously functioning pump and a custom-made 1L overflow chamber (Belleville SA, France) heated with a hot plate. Pre-programmed pumps ensured periodic injection of liquids (substrate, buffer or inocula) into the overflow chambers. A security system kept the liquid level in the reactor below a threshold. A solenoid valve (SMC, Japan) governed by a pH meter controlled the injection of CO_2_ gas to the cathode chamber. Two electronic barometers (VEGA Americas, USA) continuously monitored the gas pressure in the AC and the CC, which were kept below 5 and 10 mbar, respectively. All transmitters were purchased from Ardetem, France. The AC and the CC enclosed a reference electrode (ProSense, Netherlands) each, inserted in between the anodes and the cathode at about 1 cm from the electrodes. These reference electrodes allowed the measurement of the potential of the anodes and the cathode as well as the Ohmic loss, which was the open circuit voltage between them. These reference electrodes did not participate in the potential difference control. A multichannel potentiostat (VSP model) equipped with a current booster (Bio-Logic Instruments, France) was used for voltage control, current measurement and continuous recording of electrode potential of the TRL4 reactor. In this configuration, we have the relationship in Eq. (1), in which all values were measured by the potentiostat:

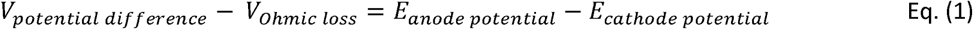

### 2.2 Bioanode Performance Modeling

All potential measurements were made against Ag/AgCl/0.6 mol kg^-1^ KCl (0.25 V vs standard hydrogen electrode) unless otherwise indicated. The two faces of the electrode were considered as being electrochemically active for calculating the current density. COMSOL Multiphysics ^®^ was used to provide model of the cyclic voltammetry (CV) of a fully developed and an “aged” bioanode. The dimension studied was 300 × 120 × 335 mm, corresponding to the reactor’s hydraulic volume. Modified Nerst-Monod equation[40] used in the model was fitted from experimental CV data. CV of the fully developed bioanode was obtained on a small (4 cm × 4 cm) carbon tissue producing 10.5 A m^-2^ current density. CV of the “aged” bioanode with decreased activity was obtained on the same carbon tissue when its activity decreased by 43 % (producing 6 A m^-2^) under exactly the same conditions (Figure S1). A multichannel potentiostat (VMP3 model) (Bio-Logic Instruments, France) was used in CV measurements with a sweep rate at 0.001 V s^-1^. These CV data was mathematically adapted and used to fit modified Nernst-Monod equations applied during the modeling (Table 1).

**Table 1.** Equation and parameters used for performance modeling of the active bioanode and the “aged” bioanode where j = current density (A m^-2^); j_max_ = maximum current density (A m^-2^); R = ideal gas constant (8.3145 J mol^-1^ K^-1^); e = the base of the natural logarithm; F = Faraday constant (96 485 Coulomb per mol^-^ e^-^); α = coefficient introduced for slope adjustment; T = temperature (298 K); and E_0.5_ = potential at which j = 1/2 j_max_.

|  | Fully developed carbon tissue | “Aged” carbon tissue | “Aged” TRL4 reactor anodes |
| --- | --- | --- | --- |
| Equation | $J(A\ m^{-2}) = \frac{J_{max}}{1 + e^{\frac{-\alpha F(E - E_{0.5})}{RT}}}$ | | |
| $J_{max}$ | $10.5\ A\ m^{-2}$ | $6.0\ A\ m^{-2}$ | 0.65 |
| $\alpha$ | 0.363 | 0.622 | 0.622 |
| $R$ | $8.314\ J.mol^{-1}.K^{-1}$ | | |
| $T$ | 298 K | | |
| $E_{0.5}(vs\ Ag/AgCl)$ | -0.3 V | -0.35 V | -0.3 V |

The CV of granular carbon cathode was obtained from the work of Marshall, et al. [41]. As it cannot be fit with a Nernst-monod equation, a polynomial equation Eq. (2) was proposed by regression (Figure S1):

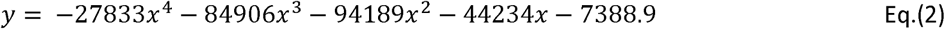

CV of the TRL4 reactor were obtained on “aged” bioanodes (day 120) with a sweep rate at 0.001 V s^-1^.

### 2.3 Experimental Setup

All chemicals were purchased from Sigma-Aldrich unless specified. During the operation, the anode in the slot close to the CEM (named slot 1) was connected with the anode in the other slot (named slot 2) with a copper cable, except when the cable was removed to measure current production of each anode. The pH of the AC was maintained at 7 by injecting 30 g L^-1^ of K_2_CO_3_ using an automatic pomp controlled by a pH meter in the anodic overflow chamber. The pH controlled CO_2_ injection to the cathode chamber was triggered when the pH of the cathodic overflow chamber exceeds 7.5. The injected CO_2_ gas came from a gas bottle (Linde, Ireland). The injection flux was about 1.5 mL per second. The anodic overflow chamber was heated to 38°C in order to have a temperature of 30°C in the AC. The cathodic overflow chamber was heated to 31°C in order to have a temperature of 25°C in the CC. The recirculation flux between the overflow chambers and the corresponding reactor chambers was 100 L per hour. Liquid samples were taken every two days for chemical analysis and conductivity measurement (Mettler Toledo, USA).

#### 2.3.1 Inoculation

Before inoculation, the AC and its overflow chamber were filled with biochemical methane potential (BMP) medium (NF EN ISO 11734) amended with sodium bicarbonate (8 g HCO_3_^-^ L^-1^) to reach a final working volume of 5.25 L in the AC and 500 mL in the overflow chamber. The CC and its overflow chamber were filled with general-acetogen-medium (pH = 7) as described in [42] to reach a final working volume of 5.25 L in the CC and 500 mL in the overflow chamber. 20 mM of 2-bromoethanesulfonic acid was added in CC for methanogen inhibition. The IC and its overflow chamber were filled with a buffer solution (35 g L^-1^ of KCl and 32.6 g L^-1^ of KH_2_PO_4_) to a final volume of 2 L in the IC and 500 mL in the overflow chamber. The anodes were inoculated with five 2 cm × 4 cm carbon-tissue-bioanodes from running H-type reactors fed with biowaste. These carbon tissues were deposited on the top edge of the anodes. The cathode inoculum was obtained from a batch reactor of dry dark co-fermentation of synthetic food waste, cardboard and centrifuged granular sludge of a mesophilic industrial UASB (upflow anaerobic sludge blanket digestion) reactor treating effluents from a sugar factory [43]. For inoculation, 100 g of the inoculum was suspended in 500 mL of distillated water and added to the CC by a pipe fitting on the lid.

#### 2.3.2 Start-up Phase

The start-up phase corresponds to the first 17 days of the operation. Sodium acetate was used as bioanode substrate during the start-up phase. At the beginning, 23.6 g of sodium acetate was added to the AC to achieve a final concentration of 50 mM in the anolyte. 23.6 g of sodium acetate was added each time when the current density fell below 0.1 A m^-2^ which indicates that all the substrate was consumed (fed-batch mode). The potential difference between the anodes and the cathode was set at 0.6 V. It was gradually increased to 1.2V with the increase of the current density.

#### 2.3.3 Biowaste Hydrolysate (BH) Feeding

At the end of the start-up phase, BH was added as substrate instead of sodium acetate. The biowaste came from an industrial deconditioning platform dealing with food waste of food industry and supermarkets. Sampled biowaste was stored at −20°C before use. Thawed biowaste was centrifuged at 6 000 g for 15 min and the supernatant was used to feed the bioanode (COD = 100 g O_2_ L^-1^). For the first ten days following the start-up phase (day 17 to day 27), 260 mL of BH was added to the AC when the current density fell below 0.1 A m^-2^, in order to have a final chemical oxygen demand (COD) around 5 g L^-1^. From day 28, an automatic feeding system was implemented, injecting 45 mL of BH (equivalent to 4.8 g of COD) per day into the AC (except for the 33^th^ day). The potential difference was 1.2 V in this phase except during the regeneration experience.

#### 2.3.4 Bioanode Activity Regeneration With Nitrogen Gas Bubbling

The potential difference was set to 0.6 V at the 40^th^ day of operation. The daily injection of BH was stopped for the 39^th^, 40^th^ and the 41^th^ day in order to analyze the regeneration effect. Nitrogen gas from a gas bottle at 3 bar pressure was injected at a flux of 670 mL per minute during 30 min to the AC via stainless steel tubes of 320 mm length and 5 mm internal diameter (Figure 2C) under the two anodes. The end of these tubes was closed up. Six apertures of 2 mm diameter were regularly perforated on the tube as the gas outlet. At the next day the potential difference was set to 0.8 V. At the 42th day the potential difference was set to 1.2 V and the daily injection of BH restarted. On day 55, the manometer of the CO_2_ gas bottle loosed, resulting in a gas flux out of control. The CC was emptied due to the gas pressure and the operation was stopped.

#### 2.3.5 Bioanode Activity Regeneration With The Double Anode Configuration

An independent assay was carried out on the reactor to study the bioanode regeneration by biofilm removal. 23.6 g of sodium acetate was added as anode substrate in fed-batch mode for the first 57 days. The potential difference was set between 0.75-1 V during this period (see Figure 3) and at 0.6 V during day 37 and 57 (Christmas holidays). From the day 57, 50 mL of BH was added in fed-batch mode in the AC. The potential difference was gradually increased to 0.9 V. After the occurrence of the aging effect, the anode in slot 1 was removed from the reactor. The stainless steel grill and the carbon tissue strips were renewed, and the frame was cleaned by water and then 70% ethanol. The anode in slot 2 was moved to the slot 1, and the cleaned anode was put in the slot 2. During this process, the reactor was always under polarization. The CC was not inoculated.

**Figure 3.**
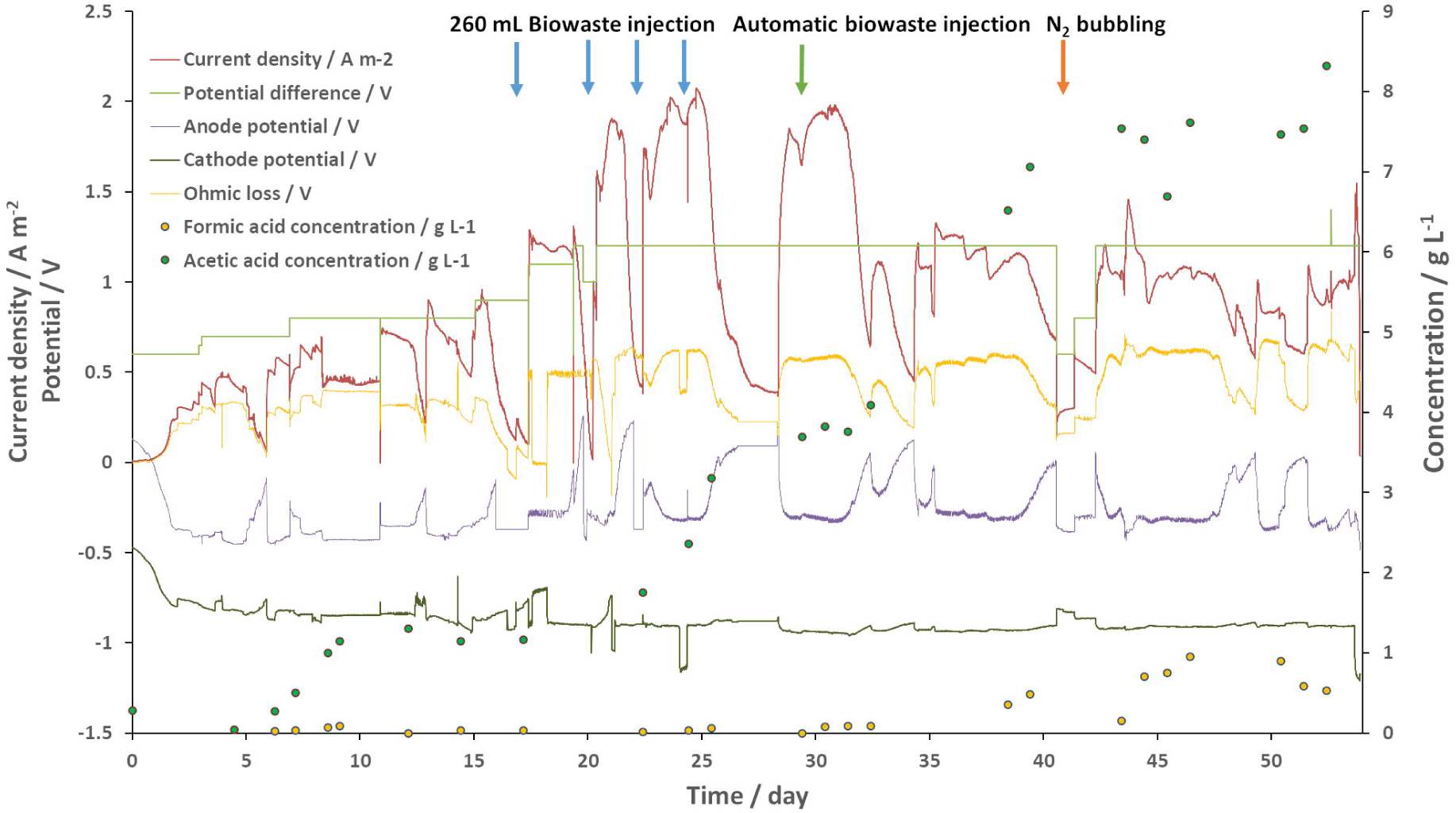
Performance synthesis of the TRL4 reactor. The y-axis on the left indicates the current density and the potential. The y-axis on the right indicates the concentration. Anode and cathode potential versus Ag/AgCl electrode were presented. Concentrations of VFAs detected in the IC are presented.

#### 2.4 Chemical Analysis

The chemical oxygen demand (COD) was measured with the LCK514 kit (Hach Lange, Germany). Volatile fatty acids (VFA) concentrations were measured with liquid chromatography (DIONEX, USA). Gas compositions (O_2_, N_2_, CH_4_, CO_2_, H_2_ and H_2_S) were determined with gas chromatography (Varian CP4900, USA).

### 2.5 Microbial Community Analysis

Bioanode samples were taken from the electrode by scratching. Liquid samples were centrifuged at 10 000 g for 20 min before removing the supernatant. Total DNA was extracted using the Powersoil™ kit (MoBio laboratories, Belgium) according to the manufacturer’s instructions. Extracted DNA concentrations were quantified with the Qubit fluorescent dye assay (Invitrogen, USA). Bacterial and archaeal 16S rRNA genes were amplified and sequenced as described in [44] with the 515F (5’-GTGCCAGCMGCCGCGGTAA-3’) and 928R (5’-CCCCGYCAATTCMTTTRAGT-3’) primers [45] and the Ion Torrent PGM™ platform (Ion Torrent, USA). The 16S rDNA tag reads were analyzed with the Galaxy supported Frogs pipeline [46]. Briefly, sequences with less than 300 pb or with error in primers were discarded. The remaining sequences were truncated to 300 bp and dereplicated. Singletons were removed. Operational Taxonomic Units (OTUs) were clustered with 97% identity. Chimera were removed. The taxonomy affiliation of OTUs was performed with a minimum confidence of 0.85. The BIOM1 format OTU table was analyzed and visualized by the Shiny Easy 16S interface (http://genome.jouy.inra.fr/shiny/easy16S/)[47]. Briefly, alpha rarefaction, Shannon index and Bray-Curtis dissimilarly was calculated using the phyloseq R package[48].

## 3 Results And Discussion

### 3.1 Pilot operation and bioanode activity regeneration with nitrogen gas bubbling

The TRL4 reactor had a total volume of 12.5 L ± 1 L, depending on the filling level. If the overflow chambers were included, this volume reached 14 L ± 1 L. The anode surface in this article was calculated with the projected surface of both electrode faces, resulting in 0.18 m^2^ per anode plate. The surface over volume ratio of the AC was 68.6 m^2^ m^-3^. The cathode pH raised to 7.6 during the first day and remained between 7.6 and 8.0 during the whole operation (Figure S2). The CO_2_ gas was thus injected continuously to the CC, establishing a permanent 10 mbar overpressure, with CO_2_ gas partial pressure > 8.0 mbar. The pH of the IC was very close to the pH of the CC. This effect could be explained by abiotic water hydrolysis which produced OH^-^ which can balance the acidity of the carbonic acid and can cross the AEM. The conductivity evolution (Figure S2) of the anode and IC followed expected trends. As bioanode activity tends to decrease the pH [49], a 5.3 S m^-1^ K_2_CO_3_ solution was injected to the AC when its pH went below 7. The anolyte conductivity was equilibrium of ion loss due to electric force and gain from the pH buffer and substrates additions, which were both positively correlated to the bioanode activity. The conductivity of the AC slightly decreased during the start-up phase, but constantly increased to 5 S m^-1^ since the injection of BH (pH = 3.9). The conductivity of the IC received ions from both AC and CC and increased continuously from 3.7 to 9.8 S m^-1^. It decreased after the N_2_ bubbling process, probably due to the voltage decrease during this process that decreased the ion diffusion due to electric force. The conductivity of the CC also increased drastically during the operation to reach 10 S m^-1^ at the 30^th^ day. We couldn’t identify the exact cause of this phenomenon. Anions such as OH^-^, acetate and CO_3_^-^ should diffuse from the CC into the IC. With no cation entrance, the catholyte conductivity should slightly decrease during the time because of water hydrolysis while the contrary was observed. Our hypothesis was that cations entered the CC by a minor leak or through the AEM because anion exchange membranes are not completely impervious for monovalent cations.

Gas produced by the AC was analyzed at the 18^th^ day (end of the start-up phase) and the 45^th^ day. At the 18^th^ day, the gas phase contained 81% of CO_2_, 11% of CH_4_ and 0.5% of O_2_. No hydrogen was detected. At the 45^th^ day, the gas phase contained 52% of CO_2_, 11% of CH_4_ and 30% of N_2_, 0.2% of H_2_ was detected as well. Gas produced by the CC on the same days were composed by 98% of CO_2_ and 1% of H_2_, with little changes during the time. These results indicate that AC lost a part of the input substrate due to methanogenesis. In contrast, the methanogen inhibition at CC was successful.

Electrochemical data acquired during the operation were summarised in Figure 3. As neither the anode potential nor the cathode potential was fixed, the voltage and the Ohmic loss determined electrode potentials. Data presented in Figure 3 confirmed Eq. 2 at all moments of the operation. The fraction of energy wasted due to Ohmic loss can be calculated by *V*_*Ohmic loss*_ */V*_*potential difference*_ which was directly correlated to the current density. It approached to zero when the current density was low and reached 50% at the maximum current density. The cathode was designed to have a more stable potential than the anodes with a significantly larger active surface. As shown in Figure 3, the cathode potential remained at about −0.9 V vs Ag/AgCl, despite considerable variation of the anodic potential from −0.4 V to 0.2 V. Sudden variations of the cathodic potential appeared during the fed-batch operation (day 0 to day 25) due to AC K_2_CO_3_ injection triggered by substrate feeding. The saline solution could significantly lower local Ohmic loss and thus pull up the cathodic potential. This issue is discussed in the next paragraph.

The anodic potential depended more on its own activity than on the imposed potential difference. As shown in Figure 3, the anodic potential can increase from −0.4 V to 0.2 V if the substrates were used up. When the substrates were available again, the anodic potential instantly drops between a narrow zone between −0.3 V to −0.4 V vs Ag/AgCl[50], depending on the imposed potential difference. Interestingly, once this “comfort zone” reached, the anodic potential became very resistant against any change, unless the substrate was exhausted. This effect can be observed at the 16-17^th^, the 24^th^, the 28^th^ and the 40^th^ day, where the anodic potential stabilized around −0.4 V while the Ohmic loss voltage drop caused quick changes of the cathodic potential, despite the enormous active surface of the cathode compared to the anodes. The substrate availability seems thus to be the only important factor that influenced the anodic potential. The Ohmic loss voltage was positively correlated to the current density. However, after the 33^th^ day, the current density significantly decreased while the Ohmic loss voltage remained at the same level, as well as the anodic potential. If we focus on the fed-batch mode operation, a substrate injection had three simultaneous effects: (i) anode potential drop; (ii) current density raise and (iii) Ohmic loss voltage increase. Knowing that all of the three were related to the bioanode activity, the potential drop appears to be the cause while the two others would be the results. This hypothesis is supported by the potential-current relationship after 35^th^ days: the potential was very similar to the 30^th^ day potential whilst the current density decreased by 40% to 50%.

The “aging” effect appeared at the 32^th^ day. Same quantity of BH was correctly injected to the AC with all conditions similar to the day before. However, the anodic potential decreased olny at −0.2 V. The current density peak reached only 55% compared to the 31^th^ day and endured only several hours before decreasing as if the substrate was used up. Our first hypothesis was an overload effect. However, the current continued to decrease on the day 33 and did not show sign of stabilization. At day 34, the current increased with BH feed as if no available substrate existed. Chemical analysis revealed 2.13 g L^-1^ of COD and 0.16 g L^-1^ of acetate in the anolyte. Our second hypothesis was that the anodic electroactive biofilm altered, probably due to contaminants in the BH. According to the literature, populations of *Geobacter* can tightly attach to solid surfaces by conductive nanowires [51]. Therefore, shearing force could detach contaminants and expose the electroactive biofilm. Indeed, bubbling is a widely used declogging technic in water treatment facilities. The N_2_ bubbling efficiently removed anodic biofilm as indicated by the irregular anodic potential drop on the day 42 and 43. The effect was however unsustainable due to fast growth of the contaminant biofilm. Nevertheless, the current density boost together with the anodic potential pattern on the day 43 supported the hypothesis of electroactive biofilm alternation.

Globally, the development of the bioanode could be divided into three phases. (i) The growth phase, in which the current density peak increased at each feed; (ii) The fully developed phase, in which the current density output reached the maximum and (iii) The “aged” phase, in which the anodic biofilm was contaminated or altered, characterized by a sudden drop of the current density peak. Here, the maximum current produced by the double bioanode reached 2 A m^-2^. The anodic coulombic efficiency was 60.3%, 98.6% and 49.1% during the growth, fully developed and “aged” phase, respectively. The maximum COD removal rate appeared during the fully developed phase, with 0.83 gCOD day^-1^ L^-1^ (which was the value fixed for the automatic BH feeding).

Acetate and formate were produced by the biocathode, reaching a maximum concentration of 8.3 g L^-1^ and 1.0 g L^-1^ in the extraction chamber, respectively. Acetate accumulation in the IC started at day 5, but temporarily stopped at day 8, probably due to a connection issue visible on the current density curve. Impact of this event lasted until day 17 when the production restarted. The cathodic coulombic efficiency from day 0 to day 15 was 81.1%. Quick accumulation of acetate appeared during the fully developed phase, with a daily production rate of 0.53 g day^-1^ L^-1^ at a coulombic efficiency of 63.7%. As mentioned, hydrogen production was detected in this period (1 % at CC) but was not followed due to the continuous CO_2_ injection. The acetate accumulation ni the CC slowed down after the N_2_ bubbling in the AC, possibly due to the low current density during day 41 to day 43. As before, this effect lasted about 8 days until the accumulation restarted. During this period of latency, acetate concentration decreased. Therefore, it is in fact an equilibrium of production and consumption, either by cathodic acetotrophs or by leakage through the CEM. From this point of view, it would be important to maintain the current density over a certain limit, in our case the empiric value is 0.5 A m^2^.

### 3.2 CV measurements reveal complex response of bioanodes facing potential change

The modified Nernst-Monod model was a viable approximation of CV of fully developed and aged carbon tissue bianodes (Figure S1). *J*_*max*_ represents the maximum current density; the coefficient α determines the maximum slope of the curve. “Aged” bioanodes of the TRL4 reactor produced much lower current density than the carbon tissue anode, and presented a more complex CV pattern. As showed on Figure 4, the anode 1 (defined as the anode closer to the AEM) was clearly a more performant bioanode than the anode 2 (defined as the anode closer to the reactor wall). The CV of anode 1 has a distinct peak at *E* = −0.226 V beyond which the current density was reversely correlated to the anodic potential until *E* = −0.290 V. This pattern did not appear on the CV of the anode 2, suggesting functional differentiations of the anode 1 and 2 caused by their positioning. A negative peak appeared near *E* = −0.5V only on the CV of the anode 2, implying the presence of an electrotrophic biofilm. A modified Nerst-Monod function with *J*_*MAX*_ = 0.65 A m^-2^ and *E*_*0.5*_ = −0.3 V was also presented on Figure 4. Evidently, this function is not a valid model for any of the bianodes of the TRL4 reactor. Nevertheless, the maximum slope of the function which is defined by the coefficient α is similar to the maximum slope present on both CVs, appeared when E increased from −0.5V to −0.4V. On TRL4 anode CV, the slope decreased drastically when *E* > −0.4V, but never approached 0. These CV patterns suggest involvement of various electroactive communities present on each bioanode. Further studies are necessary to clarify the community evolution during the “aging” of bioanode. The difference of CV of anode 1 and 2 implied considerable differences of functional groups.

**Figure 4.**
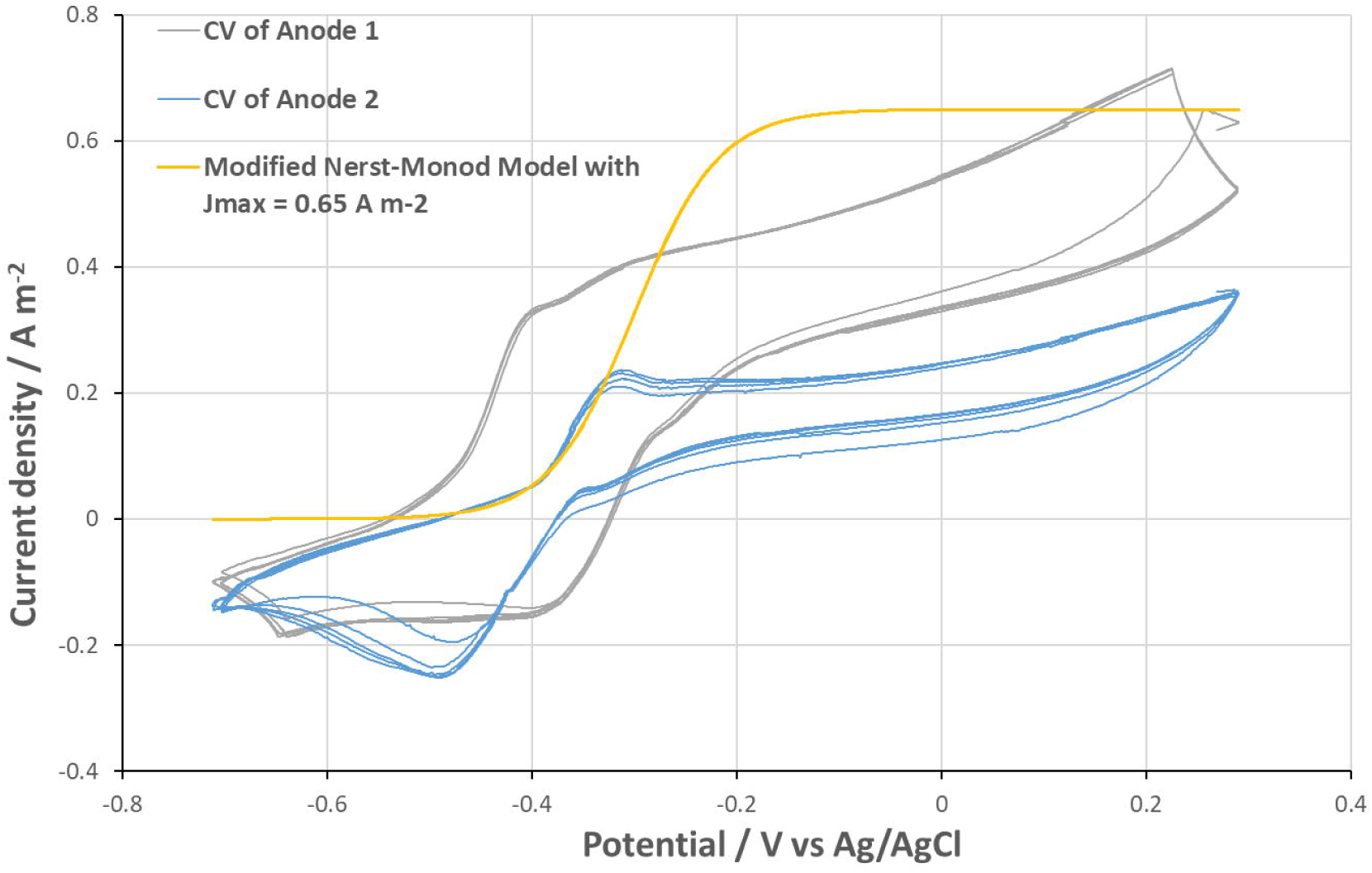
CV of aged anodes on slot 1 and 2, and a modified Nernst-Monod equation curve with ***J***_*max*_ = 0.65 A m^-2^ which was not a validate model of the CVs.

### 3.3 Bioanode activity regeneration by electrode cleaning and exchange

Figure 5 presents electrochemical data of bioanodes activity acquired during this experiment. The relationship between potential difference, anode potential, current density and Ohmic loss remained the same as discussed before. Briefly, when the substrate was available, the bioanode potential decreased to −0.4 V vs Ag/AgCl regardless of current density, potential difference or Ohmic loss. However, we can see here that an immature or an “aged” bioanode could have a less stable potential. The pattern of the cathode potential curve changed. All CC conditions (pH, CO_2_ injection, electrolyte…) were kept the same except that no bacterial inoculum was introduced. The cathodic potential was more variable under influence of the anodic potential and the Ohmic loss voltage when compared to results presented in Figure 3, while average potential was similar. This variation cannot be explained by abiotic hydrogen production, as all physical-chemical conditions were the same. The cathodic film could play an important role in potential stabilization.

**Figure 5.**
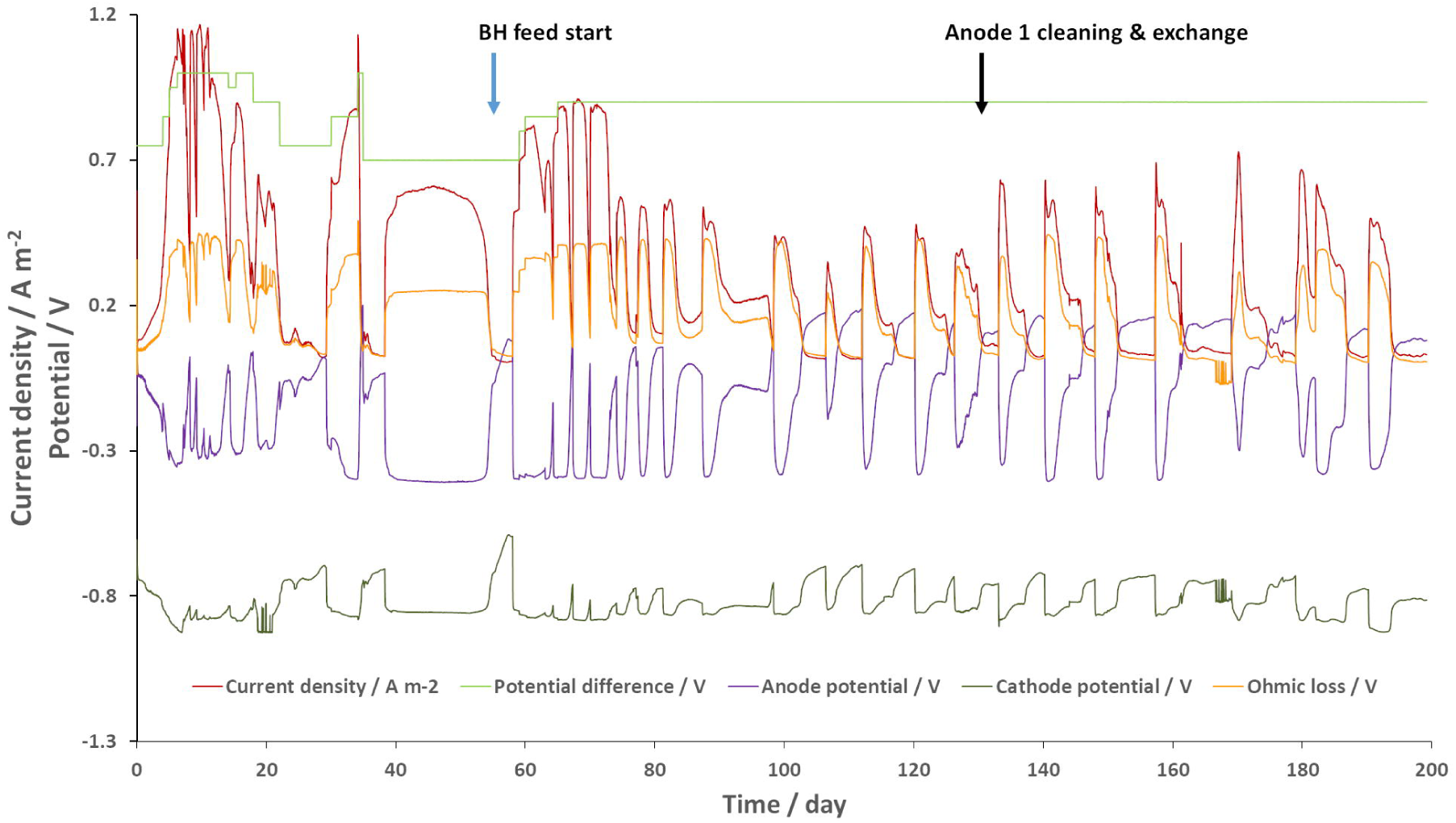
Performance of the bioanode regeneration experience. The y-axis on the left indicates the current density and the potential. Anode and cathode potential versus Ag/AgCl electrode were presented.

The current density produced by the bioanode was lower during this run as a result of lower potential difference and probably clogging of the membranes. Nevertheless, the “aging” effect appeared 14 days after the BH injection, the same as the previous operation. The current density and anodic coulombic efficiency decreases were also very similar. The “aged” bioanode was capable of producing around 50% of maximum current density of a fully developed bioanode. On the day 134, the anode 1 was cleaned and exchanged with the anode 2. In spite of the presence of an abiotic anode, the current density peak instantly increased by 30% and continued for 50 days until the next “aging” effect appeared. Average coulombic efficiency increased from 69.0% to 82.9% (average of 7 peaks before and after cleaning). Obviously, this method could not completely restore the bioanode activity. Nevertheless, it could be a way to sustain an industrial scale pilot. In addition, efficiency of the method remains to be confirmed by testing continuous operation mode.

### 3.4 Microbial community evolution

Samples analyzed by 16S sequencing were (i) inoculum of the cathode, (ii) cathode electrode and liquid phase at day 25 of the TRL4 operation, (iii) anode electrode and liquid phase before and 20-day-after the activity regeneration by electrode cleaning and exchange. The number of reads generated varied from 14 746 (cathode liquid phase) to 89 388 (Regenerated anode liquid phase), with an average of 46 484 reads per sample. The rarefaction curves (Figure S3) confirmed the representability of the analysis. A total number of 22 phyla were detected, making it unpractical to draw the classic taxonomy barplot. The Shannon index also indicated considerable alpha-diversity in all samples. As showed in Table 2, methanogens were detected in all samples despite the use of the 2-bromoethanesulfonic acid at the cathode. They were very rare on the cathode but abundant in the catholyte. This could probably be explained by the adsorption of the methanogen inhibitor by the carbon granules. For the anode, methanogens were more abundant in samples taken 20 days after the regeneration than in samples of “aged” anodes. In contrast, the electroactive *Geobacteraceae* family abundance decreased. In other words, although the electrode cleaning and exchange allowed to regenerate the bioanode activity and the coulombic efficiency, it did not happen the way we expected. We hypothesized that Geobacters would be able to colonize the clean and polarized surface before other microorganisms. The results proved that the bioanode mechanism was far more complex and Geobacter abundance is not an indicator for bioanode activity.

**Table 2.** An overview of the sequencing results. Sampling location was indicated with “top” (top of the electrode), “bottom” (bottom of the electrode” or “liquid” (electrolyte sample). “Aged” or regenerated anode samples were indicated with “Aged” or “Reg”.

|  | Number of reads | Shannon Alpha diversity | Methanogen abundance / % | Most abundant family |
| --- | --- | --- | --- | --- |
| Ino_cathode | 55122 | 1.60 | 1.33 | *Pseudomonadaceae* (37.5%) |
| Cathode | 48945 | 1.09 | 0.03 | *Pseudomonadaceae* (64.9%) |
| Cathode_liquid | 14746 | 2.64 | 23.50 | *Dysgonomonadaceae* (35.5%) |
| Aged_anode_bottom | 40166 | 2.13 | 1.34 | *Geobacteraceae* (35.6%) |
| Aged_anode_top | 32495 | 2.15 | 1.23 | *Lentimicrobiaceae* (33.1%) |
| Reg_anode_bottom | 42708 | 2.99 | 10.40 | *Dysgonomonadaceae* (10.4%) |
| Reg_anode_top | 73165 | 3.10 | 12.60 | *Dysgonomonadaceae* (16.1%) |
| Aged_anode_liquid | 21627 | 1.98 | 4.38 | *Dysgonomonadaceae* (35.9%) |
| Reg_anode_liquid | 89388 | 2.80 | 6.50 | *Synergistaceae* (17.9%) |

Figure 6 presents the Bray-Curtis dissimilarity generated with the multidimensional scaling (MDS) method. Anode samples were concentrated in the top-left zone. The regeneration process changed the microbial community on the cleaned electrode as mentioned, but not in the anolyte. The cathode community was similar to the inoculum. The catholyte had a distinct microbiota.

**Figure 6.**
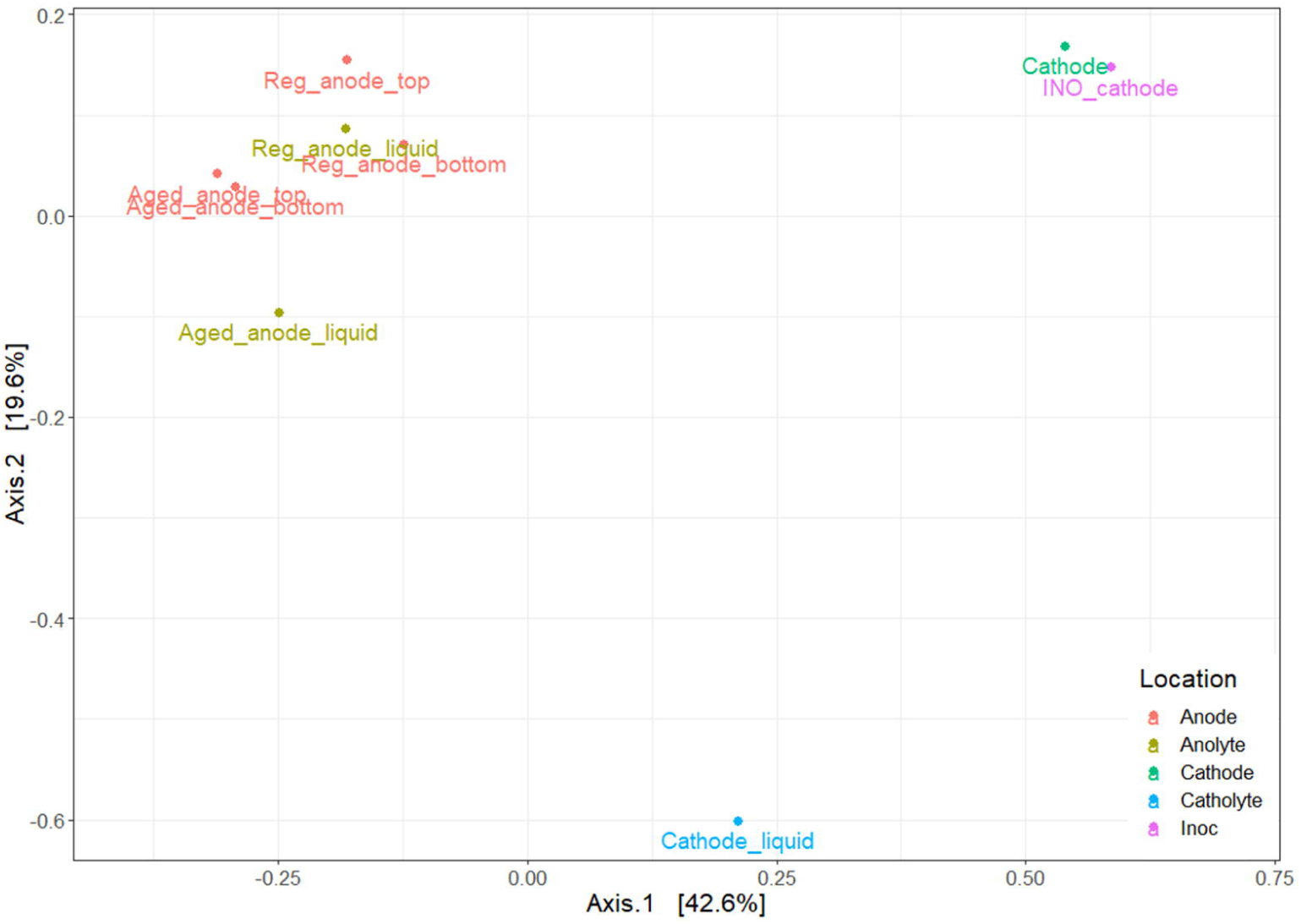
Bray-Curtis dissimilarity of sequenced sample. Sampling location was indicated with “top” (top of the electrode), “bottom” (bottom of the electrode” or “liquid” (electrolyte sample). “Aged” or regenerated anode samples were indicated with “Aged” or “Reg”.

Further analysis of the OTU table gave information of interesting but unexplained results. *Lentimicrobiaceae* and *Dysgonomonadaceae* were dominant families in all anodic samples, presenting between 10.4-35.9% of total population. The dominant genus of the *Lentimicrobiaceae* family was *Lentimicrobium*, isolated in 2016 from methanogenic granular sludge in a full-scale mesophilic upflow anaerobic sludge blanket reactor treating high-strength starch-based organic wastewater [52]. It was known for fermentative activity producing acetate, malate, propionate, formate and hydrogen. The *Dysgonomonadaceae* contained three dominant genera, *Fermentimonas, Petrimonas* and *Proteiniphilum*, all known for fermentative activities producing acetate [53, 54]. The *Synergistaceae* family (dominant in anolyte samples) was also involved in the acetogenesis [55]. These bacteria could support the bioanode activity by degrading large organic molecules into small ones such as acetate or propionate, which are common substrate for *Geobacters*. They could thus occupy the same ecological niche in the anode chamber. On the electrosynthetic biocathode, the significant dominance of the *Pseudomonadaceae* family was surprising. In this family, the only genus detected was *Pseudomonas*. Its abundance increased to 64.9% on the biocathode immerged in a mineral medium supplied only with CO_2_. However, no significant evidence supports the MES activity of *Pseudomonas* strains, to the best of our knowledge. Clearly, the abundance increase could be due to the death of other electrotrophic bacteria, but the cathode sample was taken at the day 25 (see Figure 3) when the cathodic MES had just entered the most active phase. More repetitions are still necessary to confirm this observation.

## 4 Conclusion and perspective

This work showed the feasibility at laboratory pilot scale of the concept of microbial electrosynthesis fueled by a microbial electrolysis bioanode. This concept permitted the MES to perform only with a voltage of 0.6-1.2V, which is a huge advantage compared to existing MES technologies. In the meantime, it allows the consumption of residual organic matters as a MFC or MEC do. Two experiences were carried out, revealing particular evolution of the bioanode and biocathode potentials, which could give insights of the activity the electroactive biofilms. The aging effect of bioanode was discussed as a new challenge for future application of the METs. Microbial community analysis showed a highly diversified bioanode and a specific biocathode dominated by Pseudomonas, a group of bacteria never recorded for MES.

As a first attempt of up-scaling of the novel technology, we were aware of the technical and conceptual difficulties remain to be solved. First of all, the biocathode was fragile. Its activity required a minimum electron current, which could be sometimes challenging with a bioanode. Maximizing the anodic surface may be a simple solution for this issue. Punctual current deficit could also be compensated by hydrogen gas, which could be produced and stored when there was excess current. Electrotrophic biofilm maintenance is major challenge for all MES technologies. As mentioned, the cathodic biofilm trends to disappear in long-term operations. For this issue, we cannot propose any remedy except for precise condition control and periodical re-inoculation. Secondly, the bioanode activity was highly substrate dependent. Adapted protocol would thus be necessary for each specific substrate, especially heterogeneous organic matters that could have impact on the activity of the bioanode. Finally, mechanical design is an important element of the up scaling process. We are aware of the fact that our design can be optimized in many aspects. With improved design, the anode-cathode distance and the electrode surface over volume ratio can still be ameliorated. The pumping system could also be simplified. A gas storage could be added for the CC for hydrogen and CO_2_ recovering. Solar panels can directly be used as energy source. To summarize, this technology aims not only to produce green chemistry precursors, but also to recover a large variety of residual organic matter.

## Supporting information

exactly the same conditions (Figure S1)

the whole operation (Figure S2)

The rarefaction curves (Figure S3)

## Acknowledgements

The authors of this article would like to thank Christian Duquennoi for his assistance in the establishment of the electrochemical model. The authors would also like to thank the French National Research Agency for supporting the BIORARE project (ANR-10-BTBR-02).

---

**Figure S1.** A. Cyclic voltametry of a fully matured and active bioanode (black curve) and an “aged” bioanode (red curve). The dates on the figure are not related to this work but indicate occurrence time of the “aging” effect. B. adaptation of Nerst-monod equation for kinetics of active bioanode C. adaptation of Nerst-monod equation for kinetics of “aged” bioanode. D. adaptation of a polynomial equation for kinetics of the granular carbon cathode.

**Figure S2.** pH evolution of IC and CC and conductivity evolution of AC, IC and CC during the operation.

**Figure S3.** Rarefaction curves of the sequenced sample.

