## Supplementary figures and images for "Upscaling of Microbial Electrolysis Cell Integrating Microbial Electrosynthesis: Insights, Challenges and Perspectives"

### exactly the same conditions (Figure S1)

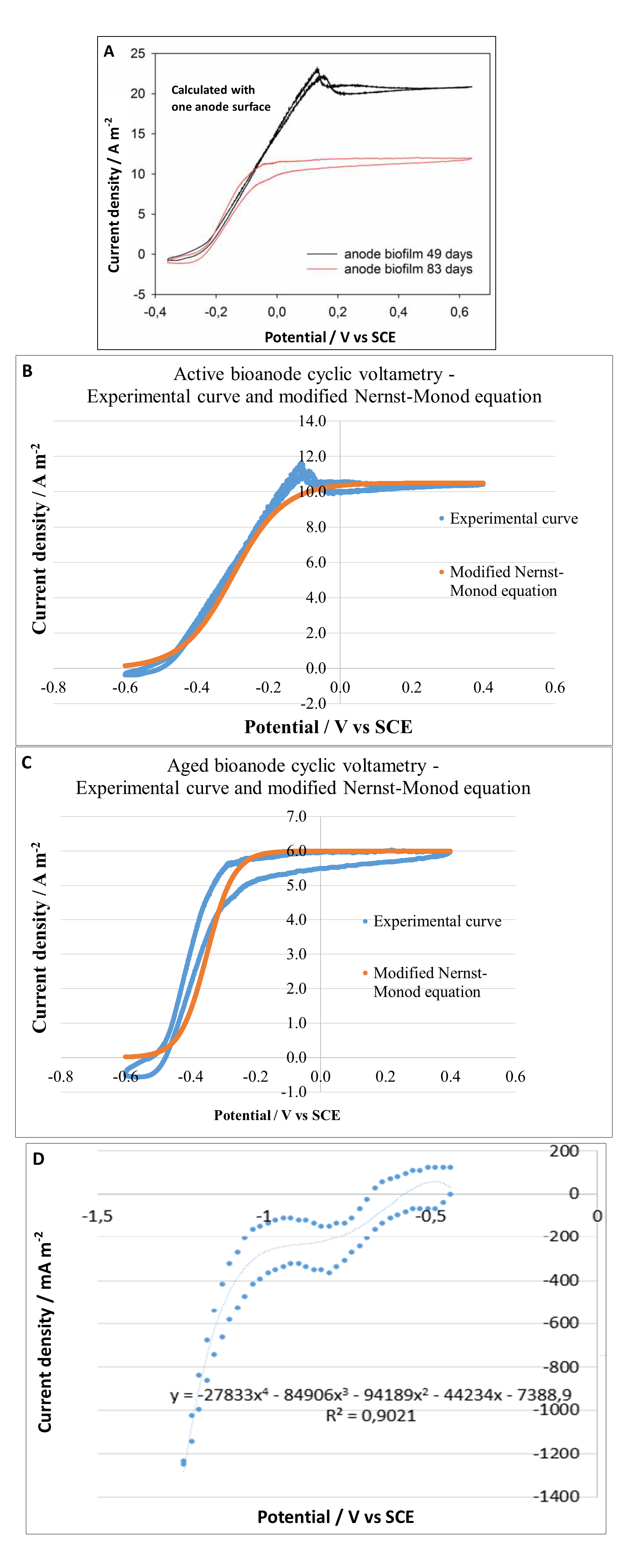

### The rarefaction curves (Figure S3)

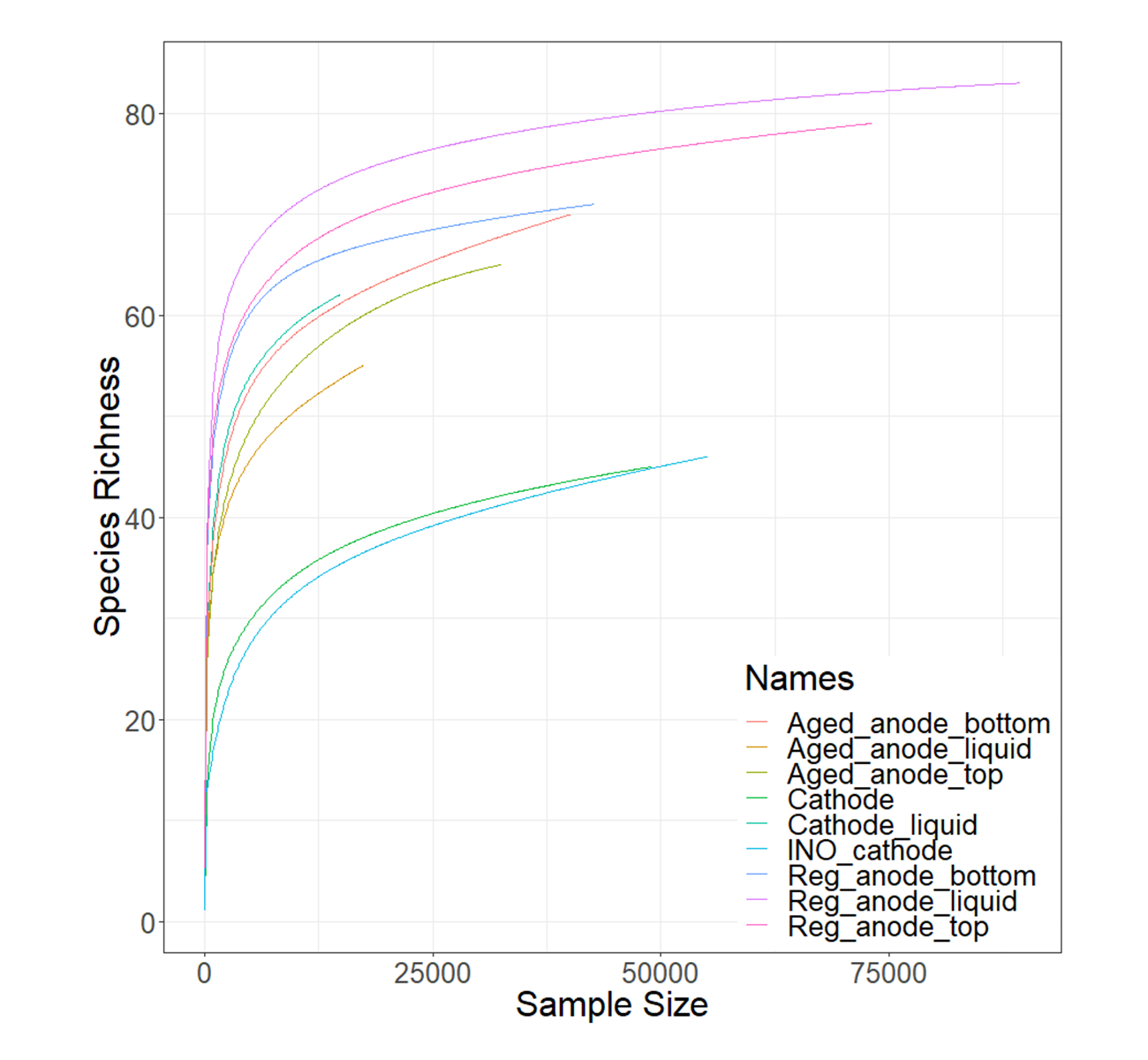

### the whole operation (Figure S2)

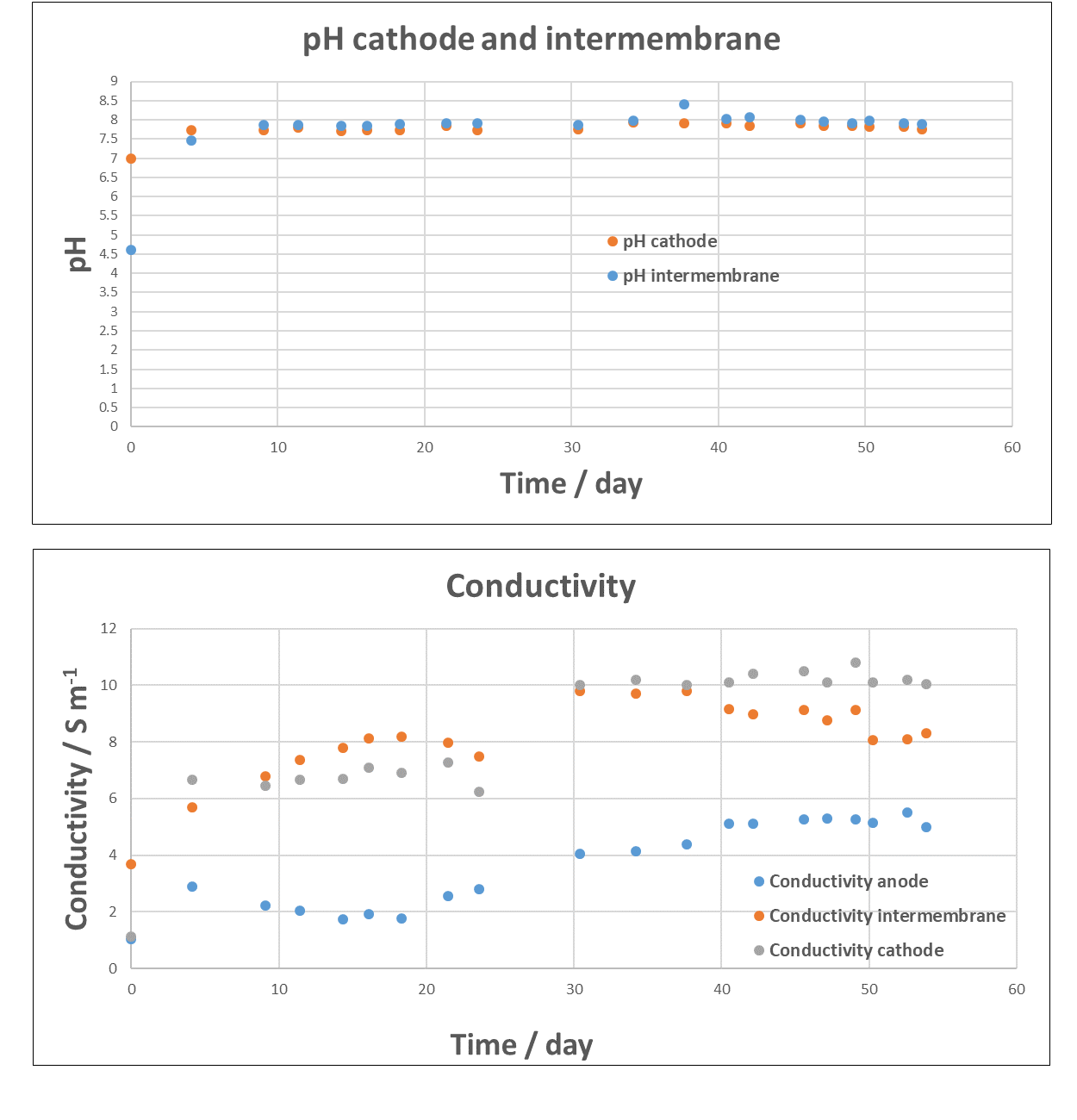
